# HIV-1 and HIV-2 differentially regulate NF-κB activity during the late stages of the replication cycle through BST-2/tetherin antagonism

**DOI:** 10.1101/2020.05.11.088385

**Authors:** François E. Dufrasne, Géraldine Dessilly, Mara Lucchetti, Kate Soumillion, Eléonore Ngyuvula, Jean Ruelle, Lionel Tafforeau, Mahamoudou Sanou, Benoit Kabamba-Mukadi

**Author notes:** Address correspondence to François E. Dufrasne.

## Abstract

HIV-2 is the second causative agent of AIDS and is commonly considered as an attenuated form of retroviral infection. Most of HIV-2-infected individuals display a slow-progressing disease, lower viral loads and a stronger immunological control of viral infection as compared with HIV-1-infected patients. The main hypothesis that could explain the difference of disease progression between HIV-1 and HIV-2 implies a more efficient T cell–mediated immunity in the control of HIV-2 infection. Herein, we investigate the effects of the HIV-2 envelope glycoprotein (Env) and its antitetherin function in the NF-κB signaling pathway during single-round infection of CD4^+^ T cells. First, we report an essential role of the Env cytoplasmic tail (CT) in the activation of this signaling pathway and we also demonstrate that the HIV-2 Env CT activates NF-κB in a TRAF6-dependent but TAK1-independent manner. Further, we show that HIV-2 reference strains and clinical isolates are unable to completely inhibit NF-κB mainly via the Env-mediated BST-2/tetherin antagonism in the late stages of the viral replication cycle in CD4^+^ T cells, in striking contrast to the HIV-1 Vpu-mediated counteraction of tetherin. We observe that this inability of HIV-2 to suppress NF-κB signaling pathway promotes stimulation of numerous genes involved in the antiviral immune response, such as *il-6, il-21* and *ifn-β* genes. Therefore, HIV-1 and HIV-2 differentially regulate the NF-κB-induced antiviral immune response mainly through the BST-2/tetherin antagonism. These new insights highlight molecular mechanisms determining, at least partly, the distinct immune control and disease outcomes of HIV-1 and HIV-2 infections.

**IMPORTANCE:** This study explores how HIV-1 and HIV-2 diverge in their regulation of the NF-κB signaling pathway. We revealed that HIV-2 fails to completely inhibit NF-κB activity, thereby inducing a stronger antiviral response than HIV-1. We demonstrated that the ability to antagonize the cellular restriction factor BST-2/tetherin largely governs the regulation of the NF-κB pathway: at the late stages of the viral replication cycle, HIV-1 Vpu blocks this pathway whereas HIV-2 Env does not. We also demonstrated that several NF-κB-targeted genes are upregulated in CD4^+^ T cells infected with HIV-2, but not with HIV-1. This stronger NF-κB-induced antiviral response may explain the better immune control of HIV-2 infection and the differences between HIV-1 and HIV-2 pathogenesis. Moreover, we observed in this study that non-pathogenic isolates of HIV-2 have an impaired NF-κB inhibitory capacity compared to pathogenic ones.

## INTRODUCTION

Tetherin (BST-2 or CD317) is an interferon (IFN)-induced restriction factor that inhibits viral release of diverse mammalian enveloped viruses by mediating their physical retention at the surface of infected cells (1–3). BST-2/tetherin poses a potent barrier to successful cross-species transmission of primate lentiviruses, such as simian immunodeficiency viruses (SIVs) to humans (1). It has been reported that an efficient viral countermeasure to BST-2 restriction was a prerequisite for the emergence of the human immunodeficiency virus type 1 (HIV-1) group M, which is responsible for the global HIV/AIDS pandemic (2). Although SIV from chimpanzee (SIVcpz), which is the immediate precursor of HIV-1, and almost all SIVs use Nef to target tetherin of their natural hosts (3–5), HIV-1 employs its accessory protein Vpu to antagonize human BST-2 (1, 6–8). This shift from Nef to Vpu occurred mainly because human BST-2 presents a deletion of five amino acids (G/DIWKK) in its cytoplasmic domain that disrupts the susceptibility to Nef (1, 9–11).

Vpu promotes surface removal, sequestration and subsequent endo-lysosomal degradation of BST-2 via a β-TrCP-dependent mechanism, reducing cell surface BST-2 levels and thereby improving the budding of virions at the sites of viral release from infected cells (12–16). In contrast to HIV-1, HIV-2 derived from SIV infecting sooty mangabey (SIVsmm) that does not encode a *vpu* gene. Interestingly, HIV-2 relies on the envelope (Env) glycoproteins to antagonize BST-2/tetherin (17), similarly to several other viruses such as Ebola virus (EBOV), Feline immunodeficiency virus (FIV), Herpes simplex viruses 1 and 2 (HSV-1/2) or Japanese encephalitis virus (JEV) (10, 17–19). BST-2 and HIV-2 Env interact through their ectodomain and the subsequent sequestration of the complex requires an endocytosis motif (GYxxΦ) in the cytoplasmic domain of HIV-2 Env (17). In addition to the aforementioned motif, several amino acids in the ectodomain of HIV-2 Env, such as T568, A598 and N659 residues are involved in ability of HIV-2 Env to counteract human BST-2/tetherin (20, 21). Interestingly, a recent study reported that these residues are highly conserved in both HIV-2 and SIVsmm strains, and that the Env proteins of some SIVsmm strains are able to antagonize human BST-2 (22). These results explain how SIVsmm has been transmitted to humans on at least nine independent cross-species transmissions and reveal a preadaptation of SIVsmm for the Env-mediated antagonism of human BST-2/tetherin (22).

Notably, HIV-2 Env-mediated antagonism removes BST-2 from cell surface but, contrary to the HIV-1 Vpu-mediated antagonism, it does not promote BST-2 degradation (15, 17). Thus, HIV-2 Env may be considered a poor antagonist of BST-2, likewise Vpu proteins of HIV-1 group N or P that demonstrate a weak activity against human BST-2 (23–25).

Besides virus retention activity of BST-2, this host restriction factor can promote innate immune response by triggering the NF-κB signaling pathway upon sensing of virus assembly and budding at the cell surface (26, 27). A dityrosine motif (Y_6_xY_8_) in the cytoplasmic domain of BST-2 is involved in this NF-κB activation mostly via the recruitment of a Syk kinase and activation of TRAF6 (28). NF-κB is a key regulator of the innate and adaptive immune responses by inducing stimulation of interferon-stimulated genes (ISGs) and expression of pro-inflammatory cytokines, type I IFNs, anti-apoptotic proteins and several potent restriction factors (including BST-2), thereby inducing an antiviral state in infected and surrounding cells which impedes viral replication and spread (29–33). Intriguingly, HIVs have selected NF-κB binding sequences in their long terminal repeat (LTR) region and thus can use this transcription factor to promote the LTR-driven viral gene transcription (33–36). Nevertheless, the virus-induced NF-κB activation also primes expression of several hundreds of genes involved in the antiviral response, which are harmful for the viral replication and spread (37, 38). HIV-1 copes with this challenge by fine-tuning the NF-κB activation during the replication cycle (26, 39). Previous studies reported that HIV-1 Nef (38, 40), Vpr (41–43) and Env (44) may activate NF-κB throughout the replication cycle, but these results remain often speculative or contradictory (38). In contrast, it is well-established that HIV-1 Vpu strongly inhibits NF-κB activity in the late stages of the replication cycle by antagonizing BST-2/tetherin, by stabilizing IκBs (inhibitors of κB) and by preventing translocation of NF-κB in the cell nucleus (39, 45). This inhibition of NF-κB activity mediated by HIV-1 Vpu is dominant and allows immune evasion and viral replication of this retrovirus (46). Recently, we reported that the HIV-2 Env can activate NF-κB in HEK293T cells on the one hand, and we also revealed that, on the other hand, the HIV-2 Env antitetherin activity was not sufficient to completely inhibit NF-κB, in striking contrast to the Vpu-mediated BST-2 antagonism (47).

To better describe the molecular mechanisms used by HIV-2 to regulate NF-κB signaling pathway, we first investigate in this study the potential involvement of accessory proteins in the regulation of NF-κB activity by co-expressing HIV-2 Env with Vif, Vpr, Vpx or Nef proteins. We observed that none of these accessory proteins can significantly modulate NF-κB activity, excepted the expression of HIV-2 Vpr that leads to a slight decrease of NF-κB activity when co-expressed with HIV-2 Env proteins. Interestingly, we also revealed using siRNA-mediated down-regulation experiments that the HIV-2 Env activates NF-κB in a TRAF6-dependent but TAK1-independent manner. Furthermore, we described the region in the envelope cytoplasmic tail (CT) involved in this activation.

In order to define the regulation of NF-κB by HIV-2 during its replication cycle and compare this to that of HIV-1, we monitored NF-κB activity in CD4^+^ T cell lines infected with HIV-1 or HIV-2 reference strains and clinical isolates. Using wild-type or *vpu*-deleted HIV-1, we confirmed that HIV-1 Vpu is a potent inhibitor of the NF-κB signaling pathway at the late stages of the viral replication cycle. Vpu exerts this role mainly through its antitetherin activity and stabilization of IκBs, as demonstrated in previous studies (27, 39). In contrast, HIV-2 Env acts as a potent activator of NF-κB in infected CD4^+^ T cells and, in the late stages of the replication cycle, HIV-2 fails to completely inhibit NF-κB activity through the Env-mediated BST-2 antagonism or any other mechanism in our experimental design. As a consequence, NF-κB activity remains important at the end of the HIV-2 replication cycle and triggers the expression of several NF-κB-targeted genes involved in the antiviral response or cell survival, as we reveal by measuring mRNA expression levels of *il-6, il-21, ifn-β* and *bcl-2* genes. This study highlights the role of NF-κB in the antiviral response and the differential regulation of this transcription factor by HIV-1 and HIV-2 viruses. Moreover, our results emphasize the importance of the viral countermeasure of BST-2/tetherin in the immune evasion capacity of HIVs. These new insights may explain, at least in part, the contrasting pathogenesis and host immune control of these two retroviruses.

## RESULTS

### HIV-2 accessory proteins do not significantly modulate the NF-κB activity

In a previous paper, we showed that the HIV-2 envelope glycoprotein activates the NF-κB signaling pathway in HEK293T cells stably expressing firefly luciferase under the control of an NF-κB-dependent promoter (47). We reproduced similar experiments but in the present study, we used HEK293T cells stably expressing a short-lived version of the firefly luciferase protein (Luc2P) that encodes a PEST sequence decreasing its stability (48). This short-lived luciferase protein is therefore a suitable model to accurately monitor activation and inhibition of NF-κB in the cells. Transduced HEK293T cells were transfected with plasmids encoding Env or Nef proteins of either HIV-1 or HIV-2. Luciferase activities were determined to assess the effect of these proteins on NF-κB activity. In agreement with published data (44), our results showed that the HIV-1 Env activates NF-κB ~ 15-to 19-fold compared to the negative control (Fig. 1A). As shown in our previous paper (47), expression of HIV-2 Env protein potently enhances the NF-κB activation with a ~ 20-to 25-fold increase compared to the negative control (Fig. 1A). These findings confirm that HIV-2 Env acts as a potent activator of the NF-κB signaling pathway in HEK293T cells.

**Fig. 1.**
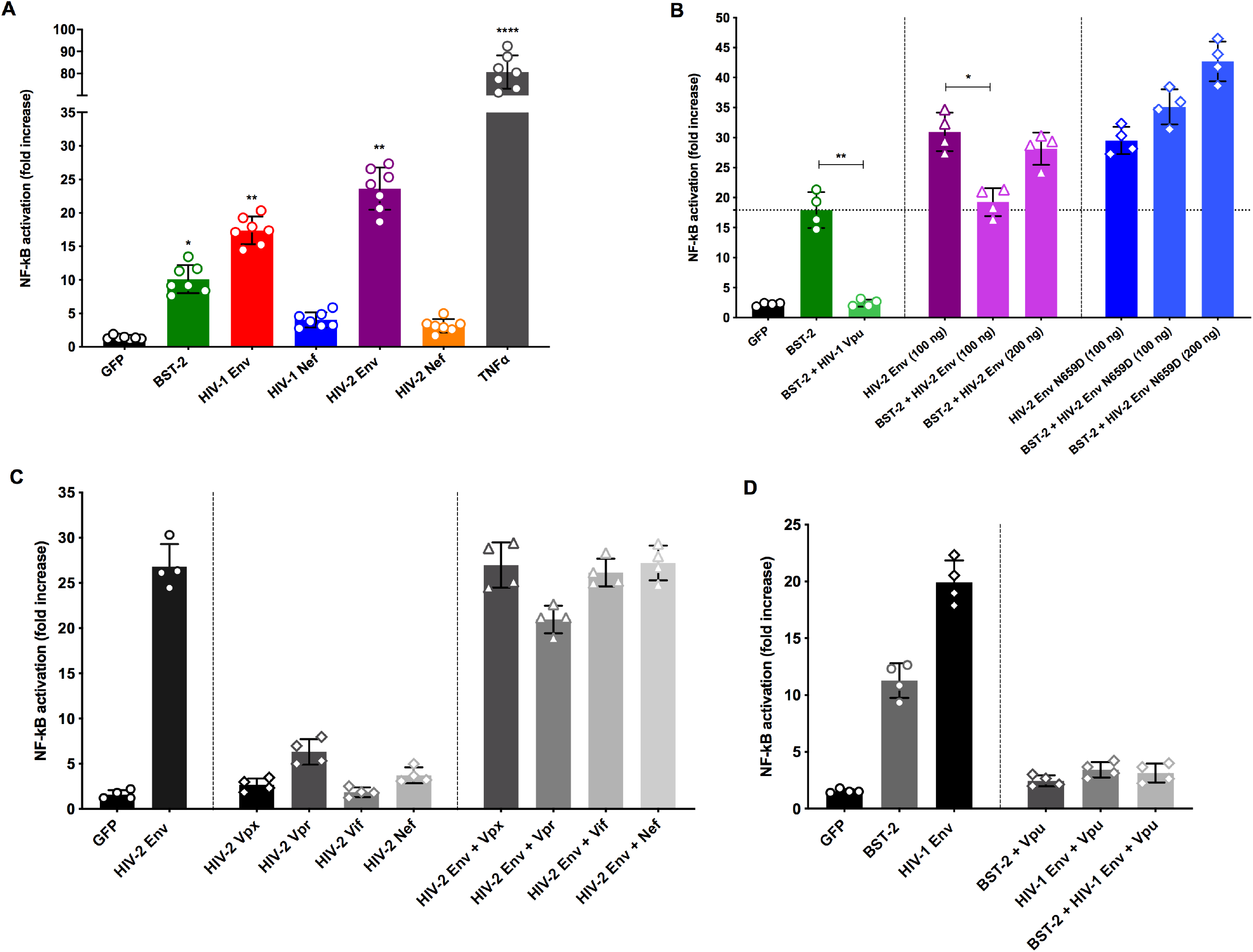
Modulation of the NF-κB activity upon expression of human BST-2, HIV-1 or HIV-2 proteins expressed alone or in combination. (A) HEK293T cells stably expressing a short-lived version of the firefly luciferase under the control of an NF-κB-dependent promoter were transfected with plasmids encoding GFP (200 ng), human BST-2 (70 ng) or viral proteins from HIV-1 or HIV-2 (i.e. Env or Nef proteins; 100 ng of DNA plasmids). HEK293T cells were lysed and luciferase activities were determined 24 hours post-transfection. As a positive control, TNFα (20 ng/ml) was added into the cell culture medium to induce NF-κB. (B) Transduced HEK293T cells were co-transfected with plasmids encoding GFP, BST-2, viral proteins of HIV-1 (Vpu; 150 ng) or HIV-2 (Env or Env N659D). Co-transfections were carried out by varying Env and Env N659D plasmid amounts (100 ng or 200 ng) while maintaining constant BST-2 plasmid amounts (70 ng). Luciferase activities were determined 36 hours post-transfection. (C) Transduced HEK293T cells were either transfected with expression vectors encoding GFP, HIV-2 accessory proteins (Vpx, Vpr, Vif or Nef; 150 ng) or co-transfected with HIV-2 Env (100 ng) and HIV-2 accessory protein expression vectors (150 ng), as indicated. (D) Transduced HEK293T cells were transfected or co-transfected with plasmids encoding GFP, BST-2 (70 ng), HIV-1 viral proteins Env (100 ng) and/or Vpu (150 ng). All panels show NF-κB activity represented as fold increase, being the ratio between the luciferase activity (RLU) of the corresponding measure and that of transduced but non-transfected HEK293T cells. These results derivate from seven independent experiments (n = 7) in (A) and four independent experiments (n = 4) in (B), (C) and (D). Data were analyzed by one-way ANOVA test followed by a multiple comparison test that compares each mean. (*, *P* < 0.05, **, *P* < 0.01 and ****, *P* < 0.0001). Error bars indicate mean ± standard deviation (SD).

We also sought to reassess the ability of the accessory proteins Nef from HIV-1 and HIV-2 to activate NF-κB signaling pathway and we found that NF-κB activity was not significantly induced by HIV-1 or HIV-2 Nef proteins (Fig. 1A).

Human BST-2/tetherin is a restriction factor that can activate the NF-κB signaling pathway, thereby triggering potent antiviral and pro-inflammatory responses (26). Here, we confirmed that exogenous expression of BST-2/tetherin induces the NF-κB signaling pathway (Fig. 1A and B). Several studies described that HIV-1 Vpu is able to overcome BST-2 restriction by removing it from the cell surface, thereby facilitating viral release of nascent virions from infected cells. Vpu binds BST-2 through its transmembrane domain and causes the subsequent endo-lysosomal degradation of this restriction factor (14, 49, 50). In accordance with these results, we observed that the NF-κB activity was markedly abolished when HIV-1 Vpu and BST-2 were co-expressed together (Fig. 1B). Moreover, according to our previous data (47), we could confirm that co-expression of HIV-2 Env and BST-2 resulted in an incomplete inhibition of NF-κB activation in HEK293T cells, in marked contrast to the potent Vpu-mediated inhibition (Fig. 1B). We also observed that increasing expression of HIV-2 Env proteins over BST-2 results in an enhanced NF-κB activation (Fig. 1B).

We showed in a previous study that the N659D substitution in the extracellular domain of the HIV-2 Env protein, as well as a truncated cytoplasmic tail, impaired the Env-mediated antagonism of BST-2. We also demonstrated that the HIV-2 Env N659D mutant proteins had lost the ability to bind BST-2 (20). Thus, we used this HIV-2 Env mutant in order to determine whether the inability of counteracting BST-2/tetherin may produce an increase or decrease of the NF-κB activity. We observed that the HIV-2 Env N659D proteins activate NF-κB in a similar way to HIV-2 Env wild-type when expressed alone. Nevertheless, in the presence of BST-2, the resulting NF-κB activation was not reduced (Fig. 1B). On the contrary, NF-κB activity was increased. Therefore, the lack of antagonism of BST-2 by Env N659D proteins results in a stronger activation of NF-κB than the antagonism-competent HIV-2 Env wild-type. Indeed, in absence of Env-mediated antagonism of BST-2, it is tempting to hypothesize that both BST-2 and HIV-2 Env N659D proteins remain free to activate NF-κB.

Given that the induction of the NF-κB signaling pathway by HIV-2 accessory proteins (Vpx, Vif, Vpr and Nef) has still not been fully elucidated, we aimed to assess whether these viral proteins possess the ability to activate or regulate NF-κB activity. Intriguingly, we observed that none of the accessory proteins from HIV-2 could significantly modulate the NF-κB signaling pathway. However, our results showed that the HIV-2 Vpr proteins can modestly activate NF-κB when expressed alone in HEK293T cells (Fig. 1C). Then, we sought to investigate whether these HIV-2 accessory proteins can modify the activation of NF-κB induced by expression of HIV-2 Env. Our data showed that the potent activation of NF-κB upon HIV-2 Env expression was not significantly up- or down-regulated by Vpx, Nef, Vif or Vpr proteins in HEK293T cells. Interestingly, however, we observed a weak decrease of the NF-κB activity upon co-expression of HIV-2 Env and Vpr proteins (Fig. 1C). Overall, these results indicated that all accessory proteins from HIV-2 are unable to significantly modulate the NF-κB signaling pathway, although HIV-2 Vpr demonstrated a trend to regulate this signaling pathway depending on the initial level of NF-κB activation.

Sauter *et al*. reported that Vpu can suppress NF-κB activation in HIV-1-infected cells in a dominant manner (39). In line with this previous data, we showed that NF-κB was not activated upon co-expression of HIV-1 Env and Vpu. Furthermore, we demonstrated that NF-κB was not activated when BST-2 and HIV-1 Env were co-expressed with Vpu (Fig. 1D). These findings confirm that inhibition of NF-κB activity by Vpu is dominant even in presence of BST-2 and/or HIV-1 Env.

### The HIV-2 envelope cytoplasmic tail activates NF-κB in a TRAF6- and MEKK3-dependent manner

We aimed to investigate the signaling cascade triggered by HIV-2 Env glycoproteins to activate NF-κB. To this end, we generated several siRNA-based knockdowns of proteins typically involved in the NF-κB signaling pathway. It has been reported that BST-2/tetherin can induce NF-κB activity through activation of TRAF6 and subsequently TAK1 (26–28). As expected, after siRNA-based depletion of TAK1 or TRAF6 proteins, we observed that the NF-κB activity decreases significantly compared to the activation induced by BST-2 in presence of TAK1 (Fig. 2A) or TRAF6 (Fig. 2B). According to the study conducted by Postler and Desrosiers, the cytoplasmic tail of HIV-1 Env stimulates NF-κB activity through the recruitment of TAK1 (44). In correlation with these results, we confirmed that inhibition of TAK1 expression markedly reduces NF-κB activity upon HIV-1 Env expression (Fig. 2A). Furthermore, we sought to identify proteins used by HIV-2 Env to potently activate NF-κB signaling pathway. Among the panel of proteins downregulated by siRNA-based experiments, we demonstrated that NF-κB activation by HIV-2 Env was markedly reduced after inhibition of TRAF6 (Fig. 2B) or MEKK3 (Fig. 2C) proteins. This implies that HIV-2 Env activates NF-κB through TRAF6, and to a lesser extent MEKK3. The specificity and efficiency of knockdowns were verified by Western blot using the same cell lysates that those used for luciferase assays.

**Fig. 2.**
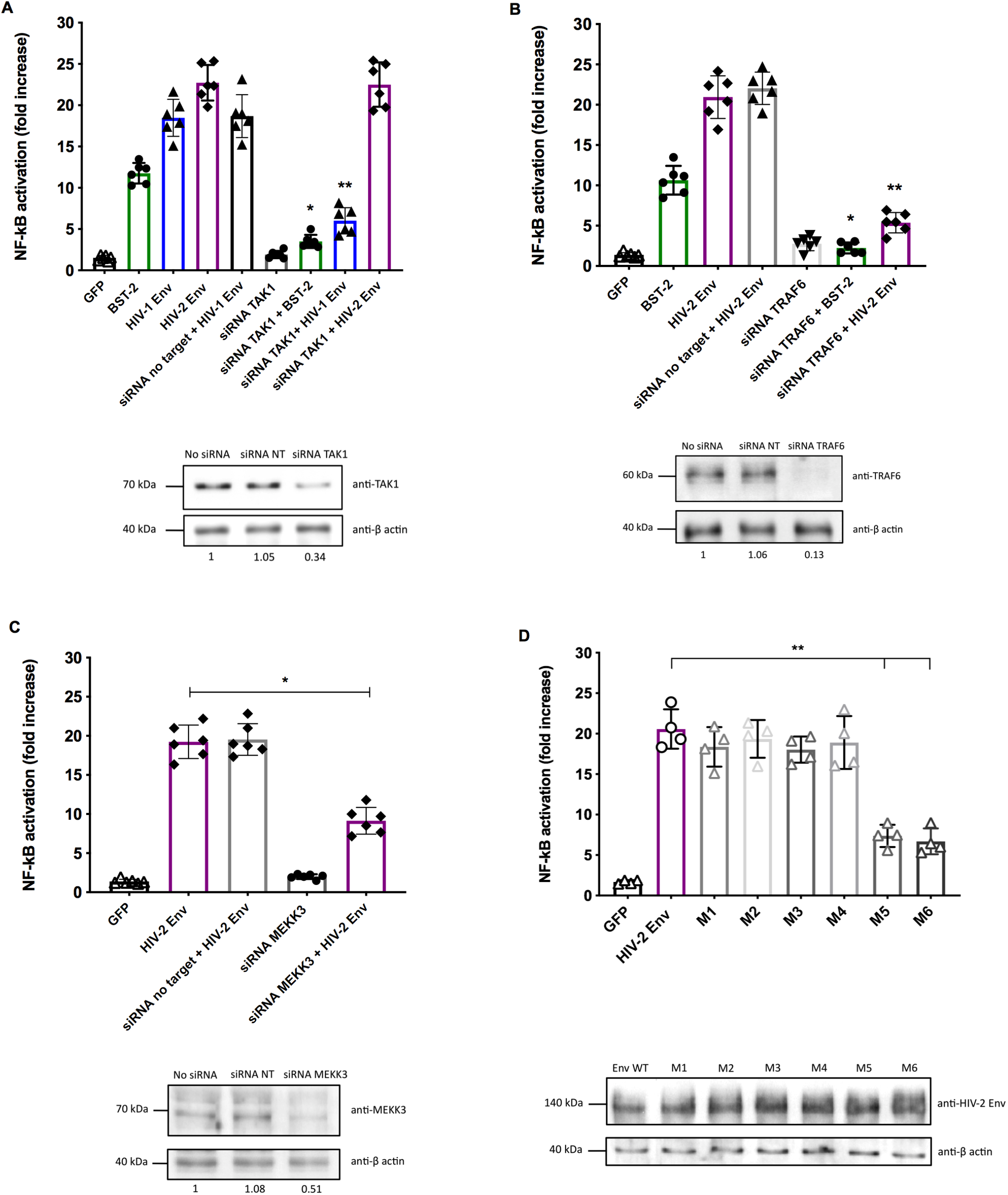
HIV-2 Env activates NF-κB through TRAF6 and MEKK3. HEK293T-NF-κB cells were either transfected with expression vectors encoding GFP (as a negative control), human BST-2 or viral Env proteins from HIV-1 or HIV-2, or co-transfected with siRNA targeting TAK1 (A), TRAF6 (B) or MEKK3 (C) mRNAs. A control siRNA (siRNA no target) was used as control. To verify the specificity of siRNA-based knockdowns, one representative Western blot is depicted below each graph. The numbers below Western blots correspond to the TAK1/β-actin, TRAF6/β-actin or MEKK3/β-actin ratios obtained after band densities determination. (D) Plasmids (100 ng) encoding wild-type Env or one of the six different HIV-2 Env CT mutants were transfected into HEK293T cells and luciferase assays were performed 24 hours later. These results derivate from six independent experiments (n = 6) in (A), (B) and (C), and four independent experiments in (D). Data were analyzed by one-way ANOVA test followed by a multiple comparison test (*, *P* < 0.05 and **, *P* < 0.01). Error bars indicate mean ± standard deviation (SD).

Postler and Desrosiers also reported that the motif YHRL in position 764 within HIV-1 Env cytoplasmic tail (CT) is largely involved in the NF-κB activation (44). Using site-directed mutagenesis experiments, we therefore built several plasmids expressing HIV-2 Env proteins truncated in the CT domain at different position. We created six different mutants (called M1 to M6) of HIV-2 Env CT and we tested their ability to activate NF-κB in HEK293T cells. Interestingly, while mutant M4 which is truncated in position 778 was still able to activate NF-κB, mutant M5 (truncated in position 758) has lost this ability. Therefore, this result strongly suggests that the region comprises between residue 758 and residue 778 within the HIV-2 Env CT is essential in NF-κB activation in HEK293T cells (Fig. 2D).

### HIV-2 fails to completely inhibit NF-κB activity mainly through the BST-2/tetherin antagonism

CD4^+^ T lymphocytes are the main targets of HIVs. In order to strengthen our previous results, we studied the modulation of NF-κB activity in CD4^+^ Jurkat cell line stably expressing the short-lived version of the firefly luciferase under the control of an NF-κB-dependent promoter. We first assessed whether our cellular model could reflect activation and inhibition of NF-κB activity over the time course. Treatment of Jurkat-NF-κB cells with TNFα induced NF-κB activity and this activation was abolished by addition of Parthenolide (PTL; 1 μg/mL), a potent inhibitor of NF-κB that directly binds to and inhibits IKKβ (51) (Fig. 3A). Then, we aimed to evaluate the NF-κB activation induced by VSV-G-pseudotyped wild-type or *vpu*-defective HIV-1 in CD4^+^ Jurkat cells. To this end, Jurkat-NF-κB cells were infected with those viruses and we found that HIV-1 completely abolished NF-κB activity at 36 and 48 hours post-infection, mainly through the Vpu-mediated BST-2/tetherin antagonism and stabilization of IκBs, as previously reported (39). Conversely, HIV-1 lacking Vpu had lost the ability to inhibit NF-κB activity compared to wild-type HIV-1 (Fig. 3B). This confirms that Vpu is a potent inhibitor of the NF-κB signaling pathway during the late stages of the replication cycle in CD4^+^ T cells (26, 27, 39).

**Fig. 3.**
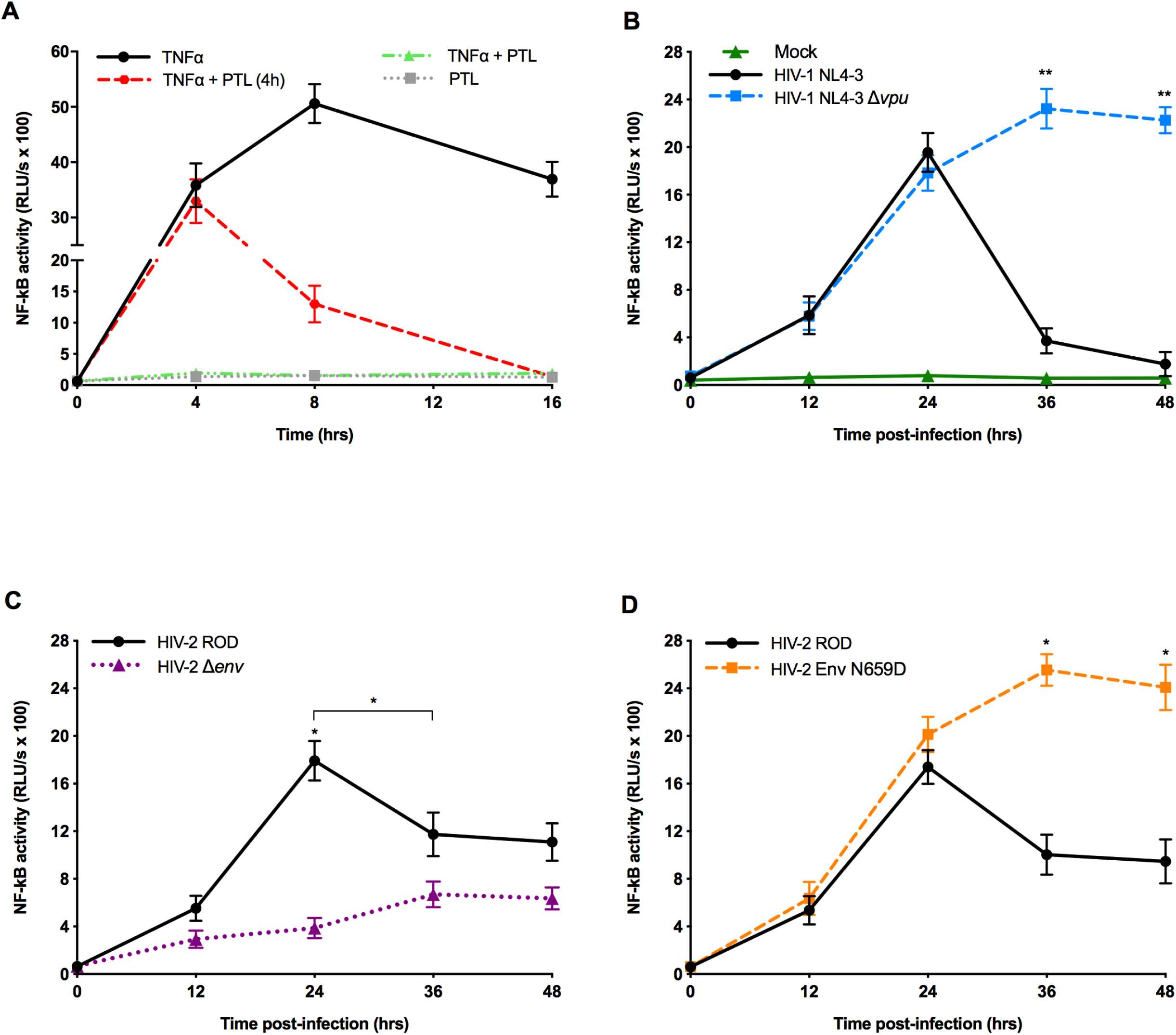
Modulation of the NF-κB signaling pathway by HIVs in CD4^+^ Jurkat cells. (A) Jurkat-NF-κB cells stably expressing a short-lived version of the firefly luciferase under the control of an NF-κB-dependent promoter were treated either with TNFα (2 ng/mL; black line) or with a specific inhibitor of NF-κB (Parthenolide, PTL; grey line), or treated simultaneously with TNFα and PTL (green line) or by addition of PTL four hours after TNFα treatment (red line). Luciferase activities were determined at different time points (4, 8 and 16 hours) after treatment. Jurkat-NF-κB cells were infected with VSV-G-pseudotyped wild-type or *vpu*-defective HIV-1 in (B), with *env-* defective HIV-2 in (C) or with HIV-2 Env N659D mutant viruses unable to interact with BST-2 in (D). Jurkat-NF-κB cells were also infected with VSV-G-pseudotyped wild-type HIV-2 ROD in (C) and (D). At 12 hours post-infection, CCR5 and CXCR4 antagonists (inhibitors of HIV entry) were added to avoid multiple rounds of infection. Luciferase activities were determined at different time points (12, 24, 36 and 48 hours) post-infection. These results derivate from four independent experiments (n = 4). Data were analyzed by one-way ANOVA test (*, *P* < 0.05 and **, *P* < 0.01). Error bars indicate mean ± standard deviation (SD).

Infection of Jurkat-NF-κB cells with VSV-G-pseudotyped wild-type HIV-2 induced NF-κB activity at 24 hours post-infection, likewise HIV-1. However and interestingly, this activation is followed by a significant but incomplete inhibition of NF-κB at 36 and 48 hours post-infection, in striking contrast to HIV-1 (compare Fig. 3B and C). By using pseudotyped wild-type or *env*-defective HIV-2, we observed that envelope trimers expressed at the surface of virions were not significantly involved in the NF-κB activation in the early stages of infection (Fig. 3C). Nonetheless, by comparing NF-κB activation induced by wild-type or *env*-defective HIV-2 viruses, we could confirm that the envelope glycoproteins are clearly involved in the NF-κB activation at 24 and 36 hours post-infection (Fig. 3C). Indeed, VSV-G-pseudotyped HIV-2 *Denv* viruses were able to infect cells but were unable to express *de novo* envelope glycoproteins in the late stages the viral replication cycle. Indeed, we observed that infection of Jurkat-NF-κB cells with these *env*-depleted virions did not induce NF-κB activation in the late stages of the replicative cycle, thereby confirming that HIV-2 Env acts as an potent activator of the NF-κB signaling pathway in CD4^+^ T cells. Moreover, these results confirmed that wild-type HIV-2 virus is unable to completely inhibit the NF-κB signaling pathway during its replication cycle, in marked contrast to HIV-1.

Next, we sought to determine whether the HIV-2 Env-mediated BST-2/tetherin antagonism is involved in the incomplete inhibition of NF-κB activity upon HIV-2 infection. To this end, we used the HIV-2 Env N659D mutant virus unable to bind and antagonize BST-2, as we reported in a previous study (20). Noticeably, infection of Jurkat-NF-κB cells with this HIV-2 mutant virus induced a stronger activation of NF-κB at 36 and 48 hours post-infection compared to that of wild-type HIV-2 virus, highlighting the importance of overcoming BST-2 restriction in the regulation of NF-κB signaling pathway during HIV infection (Fig. 3D).

In order to confirm this observation, we generated a cellular clone of Jurkat-NF-κB stably expressing specific shRNA in order to disrupt expression of human BST-2/tetherin. We used this cellular clone to assess whether wild-type HIV-2 or mutant viruses can regulate the NF-κB signaling pathway in the absence of BST-2, and we also used a Jurkat-NF-κB cellular clone transduced with shRNA no target as a control. As shown in the Fig. 4, the absence of BST-2 resulted in the inability of the HIV-2 Env-mediated antagonism to reduce the NF-κB activity. Indeed, it is conceivable that the HIV-2 Env proteins expressed at the plasma membrane remain free and available to activate NF-κB in the absence of BST-2 (Fig. 4C and D). In the case of HIV-1, the absence of BST-2 facilitates the Vpu-mediated inhibition of NF-κB activity (Fig. 4A and B).

**Fig. 4.**
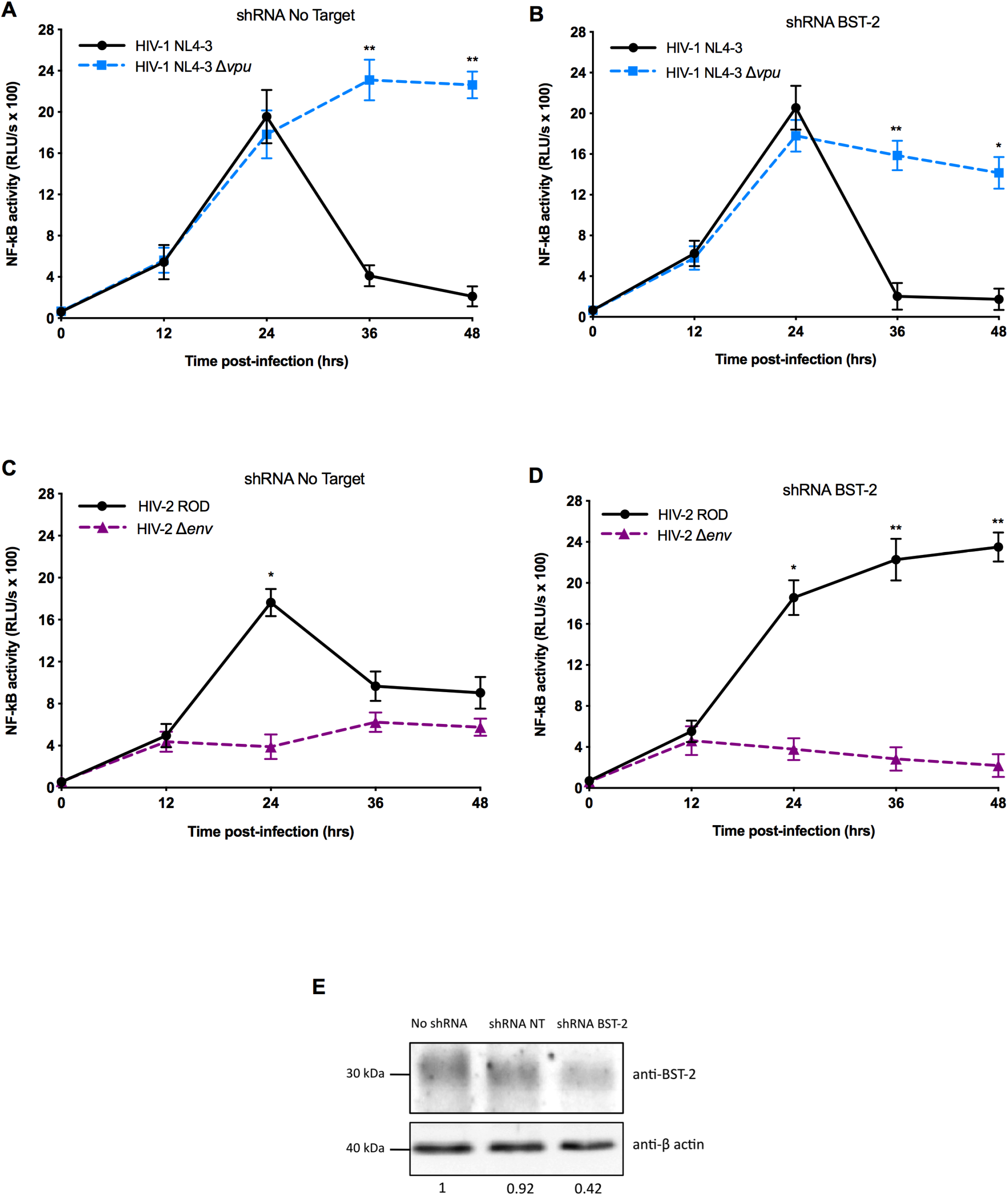
HIVs regulate the NF-κB signaling pathway mainly through the BST-2/tetherin antagonism. Jurkat-NF-κB cells were transduced with lentiviruses expressing shRNA against BST-2. Then, cells were infected with VSV-G-pseudotyped wild-type or *vpu*-defective HIV-1 in (A) and (B), with *env*-defective HIV-2 ROD or with HIV-2 Env N659D unable to interact with BST-2 in (C) and (D). Jurkat-NF-κB cells were also infected with VSV-G-pseudotyped wild-type HIV-2 ROD in (C) and (D) for comparison. At 12 hours post-infection, CCR5 and CXCR4 antagonists were added. Cells were lysed and luciferase activities were determined at different time points post-infection, as indicated in graphs. Panel (E) depicts one representative Western blot of the shRNA-based knockdown of BST-2 in Jurkat-NF-κB cells. The numbers below Western blot correspond to the BST-2/β-actin ratios obtained after band densities determination. All results derivate from four independent experiments (n = 4). Data were analyzed by one-way ANOVA test (*, *P* < 0.05 and **, *P* < 0.01). Error bars indicate mean ± standard deviation (SD).

Overall, these findings strongly suggest that the HIV-2 Env-mediated BST-2 antagonism may modestly compensate the NF-κB activation induced by Env glycoproteins, although it results in an incomplete inhibition of this signaling pathway. These results also demonstrate that the differential ability of antagonizing BST-2/tetherin largely governs the activation of the NF-κB signaling pathway during the replication cycle of HIVs.

### The inability of HIV-2 to completely inhibit NF-κB activity primes stimulation of genes involved in the antiviral immune response

Since HIV-1 and HIV-2 antagonize BST-2/tetherin by using distinct viral proteins (i.e. Vpu and Env, respectively) with contrasting efficiencies (15, 17), we aimed to define whether and how these different efficiencies of the viral counteraction of BST-2 could result in a differential regulation of the NF-κB-induced antiviral response upon infection of CD4^+^ T cells. We reasoned that the inability of HIV-2 to completely inhibit NF-κB signaling pathway in the late stages of the replication cycle may promote expression of proinflammatory cytokines involved in the antiviral immune responses. To test this hypothesis, we measured mRNA expression levels of several NF-κB-targeted genes at 24 and 48 hours post-infection by quantitative PCR (RT-qPCR).

We chose to follow expression levels of four distinct genes targeted by NF-κB (*il-6*, *il-21*, *ifn-β* and *bcl-2* genes) and we compared these expression levels after infection of Jurkat-NF-κB with HIV-1 or HIV-2. We observed that the inability of HIV-2 to completely inhibit NF-κB leads to an upregulation of these mRNA expression levels at 48 hours post-infection, in marked contrast with HIV-1 (Fig. 5A, C, E and G). For expression of IL-21, we also compared mRNA expression levels induced by infection of Jurkat-NF-κB with HIV-1 *Dvpu* or HIV-2 *Denv* (Fig. 5A). HIV-1 lacking Vpu was clearly unable to dampen the NF-κB-induced expression of *il-21* gene after 24 and 48 h of infection, and HIV-2 virus lacking Env induced a lower expression of *il-21* than HIV-2 wild-type virus. We confirmed that mRNA expression of these genes was specific of the NF-κB activation by comparing mRNA expression levels in infected Jurkat-NF-κB cells to those in infected Jurkat-NF-κB cells treated with Parthenolide, a specific inhibitor of NF-kB signaling pathway (Fig. 5B, D, F and H).

**Fig. 5.**
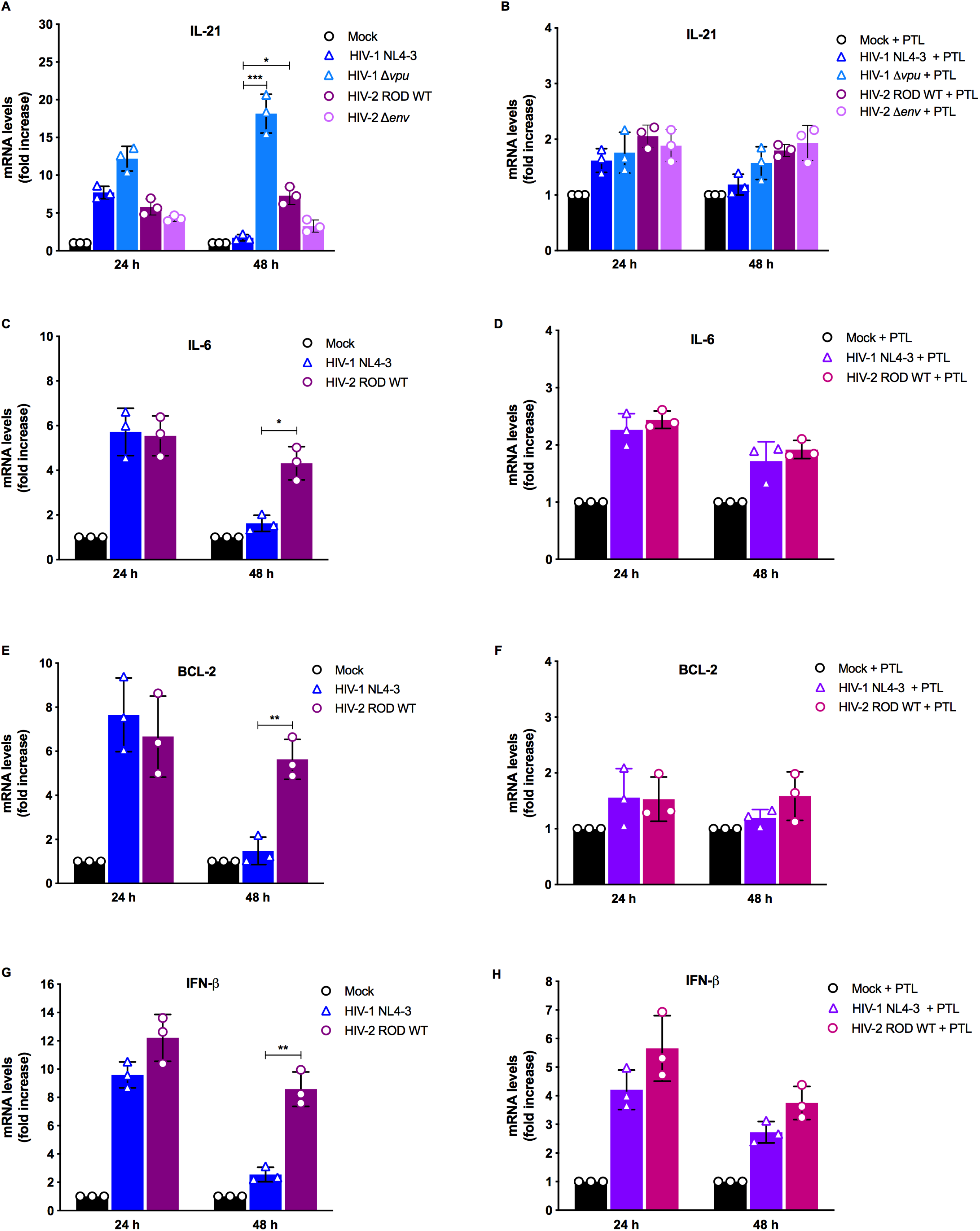
Expression levels of mRNAs induced by NF-κB during HIV-1 or HIV-2 infection. Jurkat-NF-κB cells were mock-infected or infected with VSV-G-pseudotyped wild-type HIV-1, *vpu-* defective HIV-1, wild-type or *env*-defective HIV-2 ROD and expression levels of transcripts of *il-6, il-21, bcl-2* and *ifn-β* genes (panels A, C, E, G, respectively) were measured by quantitative PCR. The ratios of each mRNA transcript were calculated relatively to those of *gapdh* at 24 and 48 hours post-infection. Panels B, D, F, H depict expression levels of mRNAs of *il-6, il-21, bcl-2* and *ifn-β* genes in infected Jurkat-NF-κB cells treated with Parthenolide. These results derivate from three independent experiments (n = 3). Data were analyzed by multiple *t* tests (*, *P* < 0.05 and **, *P* < 0.01). Error bars indicate mean ± standard deviation (SD).

### Comparison of the NF-κB regulatory capacity of HIV-1 and HIV-2 virus isolates in infected CD4^+^ T cells

In this study, we compared the NF-κB regulatory capacities of HIV-1 and HIV-2 viruses but we have only used HIV-1 and HIV-2 reference strains (lab-adapted HIV-1 NL4-3 and HIV-2 ROD strains respectively). Therefore, we aimed to confirm the differential modulation of NF-κB activity by HIV-1 and HIV-2 by using different virus isolates. Jurkat-NF-κB cells were infected by spinoculation with equivalent infectious doses of HIV-1 A018A (group M, subtype B), HIV-2 AWY (group A), HIV-2 DIL (group B) or HIV-2 7312A (group AB) virus isolates and we measured the NF-κB activity throughout their viral replication cycle. Interestingly, all HIV-2 virus isolates were unable to completely inhibit NF-κB activity at 36 or 48 hours post-infection, while infection with HIV-1 A018A virus led to a complete inhibition of NF-κB at the late stages of the viral replication cycle (Fig. 6A). Among HIV-2 isolates used in this experiment, we observed that the HIV-2 7312A (group AB) displayed the most impaired NF-κB regulatory capacity. Thus, our results suggested that the NF-κB inhibitory capacity of retroviruses may determine viral spread in the human population. Indeed, HIV-1 group M is pandemic, HIV-2 group A accounts for 80% of infection cases while HIV-2 group AB has not spread in the human population (52).

**Fig. 6.**
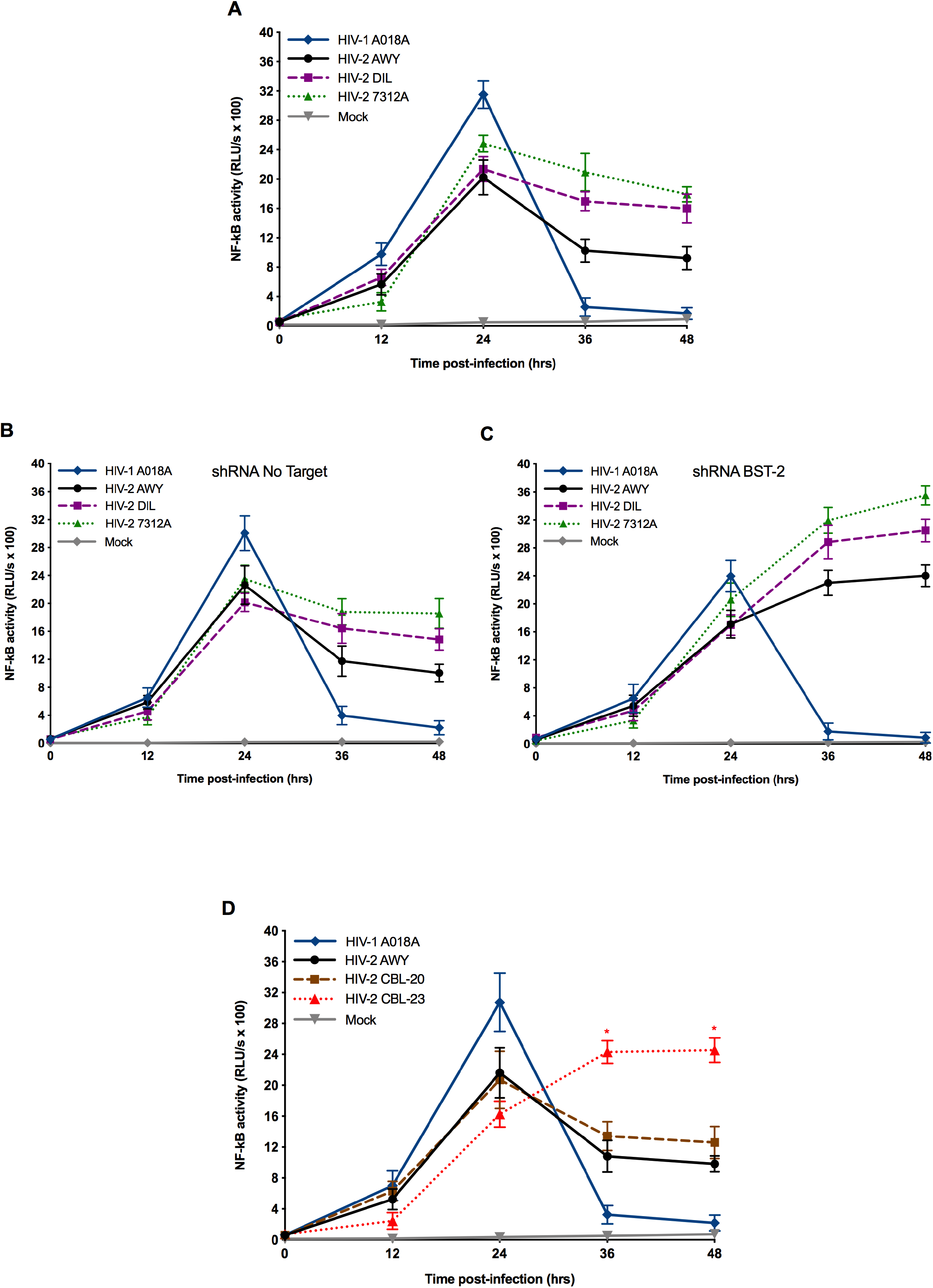
Differential modulation of NF-κB activity in Jurkat CD4^+^ T cells by HIV-1 and HIV-2 virus isolates. (A) Jurkat-NF-κB cells were mock-infected or infected by spinoculation with equivalent infectious doses of HIV-1 A018A, HIV-2 AWY, HIV-2 DIL or HIV-2 7312A isolates, and NF-κB activities were measured at 12, 24, 36 and 48 hours. NF-κB activities were also measured in Jurkat-NF-κB shRNA No Target (B) and in Jurkat-NF-κB shRNA BST-2 (C) that were mock-infected or infected by spinoculation with equivalent infectious doses of these different virus isolates. (D) NF-κB inhibitory capacities of pathogenic HIV-1 A018A, HIV-2 AWY and HIV-2 CBL-20 viruses were assessed and compared to that of non-pathogenic HIV-2 CBL-23 virus in infected Jurkat-NF-κB cells. At 12 hours post-infection, CCR5 and CXCR4 antagonists were added. Results derivate from four independent experiments in (A), (B) and (C), and from three independent experiments in (D). Data were analyzed by one-way ANOVA test (*, *P* < 0.05). In graph (D), asterisks described the statistical significance between data from HIV-2 CBL-23 and HIV-2 AWY. Error bars indicate mean ± standard deviation (SD).

Afterwards, we sought to decipher the role of the BST-2 antagonism in the regulation of NF-κB activity in CD4^+^ T cells infected with HIV-1 and HIV-2 virus isolates. As previously shown in this study, infection of Jurkat-NF-κB shRNA BST-2 cells with HIV-2 virus isolates resulted in a marked increase of NF-κB activity at the late stages of the replication cycle (Fig. 6 B and C).

Finally, we aimed to assess whether pathogenic and non-pathogenic HIV-2 virus isolates regulate the NF-κB signaling pathway. To address this issue, Jurkat-NF-κB cells were infected with pathogenic HIV-1 A018A, HIV-2 AWY or HIV-2 CBL-20 virus isolates, or with non-pathogenic HIV-2 CBL-23 viruses. NF-κB activities were monitored at 12, 24, 36 and 48 hours post-infection. Noticeably, the NF-κB inhibitory capacity of the non-pathogenic HIV-2 CBL-23 isolate was considerably impaired as compared with HIV-2 AWY and CBL-20 (Fig. 6D). Interestingly, we observed in this experiment that non-pathogenic HIV-2 CBL-23 isolate displayed an impaired ability to dampen NF-κB activity at the late stages of the replication cycle in CD4^+^ T cells than pathogenic HIV-2 isolates (Fig. 6D).

## DISCUSSION

Human BST-2/tetherin is an IFN-inducible host restriction factor that retains enveloped virions at the surface of infected cells and promotes internalization and subsequent proteasomal degradation of viral particles (6, 10, 15, 53). Besides this virus retention activity, BST-2/tetherin acts as an immune sensor of the retrovirus budding and release. Upon virus retention, a dityrosine motif (Y6xY8) in the cytoplasmic domain of BST-2 is phosphorylated and recruits a Syk kinase that can activate TRAF6 (26, 28). Ultimately, TRAF6 activates TAK1 which in turn activates NF-κB by promoting the degradation of IκBs (inhibitors of NF-κB). This transcription factor is then translocated into the cell nucleus and binds to NF-κB binding sites localized in the promoter of numerous genes that express anti-apoptotic proteins, cytokines and chemokines, IFN-β or stimulate the IFN-stimulated genes (ISGs) expression which are all able to prime an antiviral response against viral replication and dissemination (29–32). Yet, HIVs have evolved sophisticated means to dampen this antiviral immune response at the late stages of the viral replication cycle. HIV-1 Vpu proteins completely inhibit the NF-κB activity by antagonizing BST-2 (39, 54). A recent study demonstrated that the Vpu-mediated inhibition of NF-κB largely down-regulates the expression of a wide number of NF-κB-targeted genes, thereby facilitating immune evasion of HIV-1 (46).

Compared to HIV-1, HIV-2 is commonly considered as less pathogenic and is more efficiently controlled by the host immune system. It is thought that the early T cell-mediated antiviral response is a major contributor to the phenotype (55–60). In the present study, we demonstrated that the differential regulation of the NF-κB signaling pathway during the replication cycle of HIV-1 and HIV-2 may explain the differences of pathogenesis. The latter virus uses its envelope glycoprotein to antagonize BST-2/tetherin (17), but interestingly, both HIV-1 and HIV-2 Env proteins can activate NF-κB in HEK293T cells, as revealed in previous studies (44, 47). Therefore, we aimed to determine whether and how HIV-2 regulates the NF-κB signaling pathway in target cells of HIVs. Interestingly, we observed in the present study that HIV-2, in contrast to HIV-1, appeared to be devoid of a strong inhibitor of NF-κB activity and that the Env-mediated antagonism of BST-2 was less effective in decreasing the NF-κB-induced antiviral response. Indeed, in Jurkat cells infected with different HIV-2 reference strains or clinical isolates, NF-κB activity in the late stages of the viral replication cycle was three to five-fold higher than in cells infected with HIV-1.

Interestingly, we observed a dual role of the HIV-2 Vpr accessory protein, in accordance with a recent study that determined such a role of Vpr from some SIV strains (61). When expressed alone, HIV-2 Vpr slightly activated NF-κB. However, upon co-expression with HIV-2 Env, a weak inhibition of NF-κB can be observed. Therefore, it is quite probable that this inhibitory effect of HIV-2 Vpr may differ among circulating HIV-2 isolates from infected individuals. This latter issue warrants to be studied in further studies *in vivo*.

We also tested the impact of the N659D substitution within the gp36 ectodomain of HIV-2 Env, that impairs the BST-2 antagonism (20), on NF-κB activity during the viral replication cycle. We observed that the activity levels of NF-κB generated by this mutant virus were even greater than those observed during infections with the wild-type virus, suggesting that the ability to antagonize BST-2 largely governs the regulation of this signaling pathway.

It has been reported that the motif Y764HRL within the HIV-1 Env cytoplasmic tail (CT) is involved in the activation of NF-κB (44). By using site-directed mutagenesis experiments, we defined the region in the HIV-2 Env CT involved in the activation of the NF-κB signaling pathway. Truncation of the CT may affect the expression, addressing or conformation of the Env glycoproteins in a cell-specific manner (62). Yet, in HEK293T cells, functionalities of the Env glycoproteins with truncated CT are not modified (62, 63). Thus, these cells are suitable to test different envelope proteins and their ability to activate NF-κB. Although we did not define the exact residue(s) involved in the NF-κB activation, we depicted the region within HIV-2 Env CT, which corresponds to the residues localized between the position 759 and the position 777 (from HIV-2 ROD strain, GenBank: M15390.1). In this region, it is interesting to note that the HIV-2 Env CT also includes a motif YxxΦ similar to that of HIV-1 Env CT which is involved in the NF-κB activation (44). Nevertheless, further experiments using HIV-2 Env proteins harboring point mutation in this region are needed to better define the precise residues involved in the NF-κB activation. Moreover, it is still conceivable that a cellular protein may be recruited by the HIV-2 Env CT to prime the NF-κB activation. Although we have been able to demonstrate that the HIV-2 Env protein activates NF-κB via a TAK1-independent but TRAF6-dependent manner (and in a lesser extent MEKK3), the molecular mechanisms underlying this NF-κB activation remains to be fully elucidated. However and interestingly, a recent study conducted by the Guatelli’s group demonstrated that the Ebola virus matrix protein (VP40) and the envelope glycoprotein (GP) cooperate with human BST-2/tetherin to induce a strong NF-κB activity (53). Although the GP glycoprotein from Ebola virus is, like the HIV-2 Env, the viral antagonist of BST-2, the mechanism of antagonism is unclear. It does not remove BST-2 from the cell surface and there is no evidence of degradation of BST-2, like is the case of the HIV-2 Env-mediated BST-2 antagonism (64, 65). Therefore, the lack of degradation of BST-2 by the HIV-2 Env protein may result in the incomplete inhibition of NF-κB. Although we demonstrated that the interaction between HIV-2 Env and BST-2 resulted in an incomplete inhibition of NF-κB activity, we did not provide the mechanism(s) that could explain the sustained and persistent activation of NF-κB during the late stages of the replication cycle. We may hypothesize that the HIV-2 Env cytoplasmic tail and the cytoplasmic domain of BST-2, each activating NF-κB, remain “free” or “available” to activate NF-κB even if the interaction between these two proteins occurs. Further studies are needed to elucidate this issue. In the present study we have described that the *il-21* gene, as well as *il-6, bcl-2* and *ifn-β*, was markedly upregulated at 48h post-infection in CD4^+^ T cells infected with HIV-2, but not with HIV-1, and that was largely dependent to the NF-κB activity. A high expression of IL-21 can maintain and reinforce the cooperation between CD4^+^ and CD8^+^ T cell responses. Indeed, HIV-specific CD4^+^ T cells expressing IL-21 are enriched in the acute phase of infection in HIV elite controllers. IL-21 also contributes to control of HIV replication through both enhancing specific CD8^+^ T cell response and managing the regulatory CD4^+^ T cell (Treg) functions (66–69). We therefore may conclude that the upregulation of *il-21* gene expression during the HIV-2 infection that we have observed in our study is mainly due to NF-κB activation and may probably contribute to the better immunological control of HIV-2 infections. Interestingly, we also revealed an upregulation of *ifn-β* gene expression upon HIV-2 infection (but not HIV-1) at 48h post-infection. The relationship between the IFN/ISG responses and the immunopathogenesis of HIV and SIV infections is still debated until now since both deleterious and beneficial effects of IFN/ISG responses in HIV and SIV infections have been reported. However, accumulating evidences suggest that high IFN type I (IFN-I) production during the acute phase of HIV (or SIV) infections is beneficial to prevent viral replication and dissemination, to maintain healthy CD4^+^ T cell counts and to slow the progression of the disease (reviewed in (70–72)). Notably, an important study conducted with SIV revealed that the blockade of the IFN-I signaling in rhesus macaques prior to infection or during the acute phase of infection worsens the disease progression and clinical outcome (reduced antiviral gene expression, higher viral loads, accelerated decline of CD4^+^ T cells and a more rapid progression to AIDS) (73). Moreover, in humans, it has been observed that the immune cells from HIV-1 elite controllers have an increased ability to produce IFN-I compared to those of HIV-1 non-controllers (74–76) and studies have reported that the plasma levels of IFN-I in HIV-2 individuals are significantly higher than those of HIV-1, without resulting in higher levels of immune activation (55, 77). Therefore, results presented in the present study prompt us to postulate that the early and strong IFN/ISG responses induced by NF-κB during HIV-2 infection represent a major determinant of the lower levels of viral load, the stronger immune control, and may be involved in the delayed disease progression, as compared with HIV-1 infection.

Finally, we demonstrated by comparing pathogenic and non-pathogenic HIV-1 and HIV-2 isolates that the pathogenesis of retroviruses may be governed by the regulatory capacity of the NF-κB signaling pathway during the viral replication cycle. Indeed, an impaired NF-κB inhibitory capacity could be linked to a low pathogenesis of the retroviruses and to a slow disease progression of the associated infection in the human host. HIV-2 is commonly considered as less pathogenic than HIV-1 and we demonstrated in this study that HIV-2 viruses (reference strains and clinical isolates) showed an impaired NF-κB inhibitory capacity as compared with HIV-1 viruses. Furthermore, among HIV-2 viruses, a large proportion are considered as non-pathogenic in humans since more than 60% of HIV-2-infected patients are long-term non-progressors (LTNPs) (52, 78). In relation with this, two recent studies have elegantly revealed that the NFAT regulatory capacity of Nef proteins was markedly impaired HIV-1 LTNPs and elite controllers, in contrast to HIV-1 progressors (79, 80).

Overall, our study provides new insights concerning the differential regulation of the NF-κB signaling pathway during HIV-1 and HIV-2 replication cycle. In contrast to HIV-1, HIV-2 viruses fail to completely inhibit the NF-κB activation in the late stages of the viral replication cycle in CD4^+^ T cells. As a consequence, the NF-κB-induced antiviral response during HIV-2 infection is higher than during HIV-1 infection. This stronger NF-κB-induced antiviral response may partly explain the better immune control of HIV-2 and the differences between HIV-1 and HIV-2 pathogenesis. Moreover, this study also highlights the importance of an efficient viral antagonism of BST-2/tetherin in order to ensure both viral replication and immune evasion of these two closely related retroviruses.

## MATERIALS AND METHODS

### Cells, plasmids, and reagents

HEK293T and CD4^+^ Jurkat (clone E6-1) cells were obtained from the ATCC. The TZM-bl reporter cells were obtained from the NIH AIDS Reagent Program (catalog number: 8129). The HEK293T and TZM-bl cells were maintained in Dulbecco’s modified eagle medium (DMEM; Thermo Fisher Scientific) supplemented with 10% of FBS and 0.1% of gentamicin, while Jurkat cells were propagated in complete Roswell Park Memorial Institute medium (RPMI; Thermo Fisher Scientific).

The HIV-2 ROD infectious molecular clone (pKP59-HIV-2 ROD (81)), encoding a complete Env cytoplasmic tail, has been previously described (20). The HIV-1 pNL4-3 and *vpu*-deleted pNL4-3 infectious molecular clones, as well as plasmid encoding the codon-optimized *vpu* sequence were all obtained from the NIH AIDS Reagent Program (catalogue number 114, 968 and 10076 respectively). HIV-2 *env*, *nef, vif vpr* and *vpx* coding sequences were amplified with specific primers from pKP59-HIV-2 ROD plasmid and cloned into pcDNA3.1 vector (Invitrogen). Expression vectors encoding the six different mutants of HIV-2 Env cytoplasmic tail were created by introducing stop codon in different position within the cytoplasmic tail by site-directed mutagenesis using the QuickChange II Site-Directed Mutagenesis Kit (Agilent). HIV-2 Env mutants were as follows: M1: Env L843-Stop, M2: Env L824-Stop, M3: Env L795-Stop, M4: Env L777-Stop, M5: Env L758-Stop and M6: Env G739-Stop.

The expression plasmids of human BST-2, HIV-2 Env N659D, as well as infectious molecular clones of HIV-2 Δ*env* and Env N659D have been described in our previous studies (20, 47). The pHAGE NFκB-TA-LUC-UBC-GFP-W lentivector containing NF-κB binding sites upstream of the *luciferase* gene has also been previously described (82). In order to allow an accurate monitoring of NF-κB activity in CD4^+^ Jurkat cells, wild-type *luciferase* sequence was replaced by *luciferase luc2P* sequence. The corresponding luciferase protein is a short-lived version of luciferase because it harbors a PEST sequence that decreases its stability. The *luc2p* gene was amplified from the pGL4.11(*luc2P*) vector (Promega) and inserted into the pHAGE NFκB-TA-LUC-UBC-GFP-W lentivector using *NcoI/BamHI* restriction sites. All sequences of plasmid DNA were verified by nucleotide sequencing using ABI 3500 platform (Applied Biosystems).

HEK293T and CD4^+^ Jurkat cells were transduced with VSV-G-pseudotyped viruses encoding the NF-κB-Luc2P-GFP genomic cassette. Cells efficiently transduced were sorted following GFP expression using a FACSAria III cell sorter (BD Biosciences). We generated cellular clones of each cell type by limiting dilutions. After several passages, cellular clones (HEK293T-NF-κB and Jurkat-NF-κB) that exhibited minimal luciferase expression in absence of stimuli were selected, and the responsiveness of these clones was assessed by stimulation with TNFα (Miltenyi Biotec). Transfections of HEK293T cells were performed using the Transporter 5 Transfection Reagent (Polyethylenimine; Polysciences) following the manufacturer’s instructions.

For luciferase assays, the amounts of each plasmid transfected or co-transfected into the HEK293T-NF-κB cells are provided in the figure legends. The total amount of transfected plasmid DNA was kept constant in all experiments by adding pcDNA-GFP plasmids.

The inhibitor of NF-κB signaling pathway, Parthenolide, was purchased from Invivogen and was used at a final concentration of 1 μg/mL when indicated in this study. Equivalent final concentration of DMSO were added to controls. The HIV entry inhibitor bicyclam JM-2987 (used at 300 ng/mL; cat: 8128) as well as inhibitor TAK-779 (used at 100 ng/mL; cat: 4983) have been received from the NIH AIDS Reagent Program and were both added 12 hours after all infection assays.

### Virus isolates, virus production, and infections

VSV-G-pseudotyped wild-type and *vpu*-deleted HIV-1 were generated by transfecting 3.5 x 10^5^ HEK293T cells with HIV-1 pNL4-3 or *vpu*-deleted pNL4-3 molecular clones (1.5 μg) alongside with VSV-G expression plasmids (200 ng; pMD2 from addgene). VSV-G-pseudotyped wild-type, *Denv* and Env N659D HIV-2 viruses were also generated following the same protocol. Twelve hours post-transfection, culture medium was removed, and fresh medium was added in 12-well plate. Virus-containing supernatants were harvested 40 hours later and centrifuged at 1500 x g to discard cell debris. HIV viral particles were purified using the Lenti-X concentrator (Takara) and were resuspended in fresh DMEM medium supplemented with 5% of FBS. Viral infectious titers were determined by infecting 4 x 10^4^ TZM-bl reporter cells in serial dilutions in 48-well plate and we proceeded to luciferase test to assess infectious titers of these viral particles.

Jurkat-NF-κB cells were infected in duplicat with VSV-G-pseudotyped HIV-1, HIV-2 or variants thereof with equivalent infectious doses (based on TZM-bl assays).

Virus isolates HIV-1 A018A, HIV-2 AWY, HIV-2 DIL, HIV-2 7312A were obtained from the NIH AIDS Reagent Program (catalogue number 629, 12383, 3511 respectively) while the HIV-2 CBL-20 and CBL-23 virus isolates were received from the NIBSC (National Institute for Biological Standards and Control, catalogue number 0122 and 0125). Jurkat-NF-κB cells were infected with equivalent infectious doses of those isolates by spinoculation. Virus inoculum were added to culture medium containing cells and maintained at 37°C in an incubator. Four hours later, cells with virus-containing supernatant were centrifugated at 1200 x g for 2 hours at 35°C. Cells were then washed and seeded in 24-well plate. The HIV entry inhibitors bicyclam JM-2987 and TAK-779 were both added 12 hours after infection.

### Reporter gene assays and monitoring of the NF-κB activity

Transduced HEK293T-NF-κB cells were used to assess NF-κB activation upon expression of different HIV-1, HIV-2 or human proteins. Twenty-four hours (or 36h when indicated) post-transfection, cells were washed with PBS (Thermo Fisher Scientific) and lysed with 1x passive lysis buffer (Promega) for 15 min. Cell lysates were collected and centrifuged 5 min at 15000 x g, and 20 μl of lysate were added to 100 μl of Luciferase Assay Reagent II (LAR II) containing luciferin. Bioluminescent signals were measured with a GloMax 20/20 luminometer (Promega). For the monitoring of NF-κB activity in Jurkat-NF-κB cells infected with HIV-1, HIV-2, or variants thereof, one of the replicates of each condition was used to determine luciferase expression levels at 12, 24, 36 and 48 hours post-infection. Infected cells were washed twice, counted by Trypan blue exclusion method, and ~ 3 x 10^5^ infected Jurkat-NF-κB cells were lysed with 1x passive lysis buffer and NF-κB activity was measured using Dual-luciferase assay as explained above. The NF-κB activity in mock-infected Jurkat-NF-κB cells was also measured during 48 hours of infection and remained at the basal level of luciferase expression (~ 80-100 RLU/s).

### SiRNA- and shRNA-based down-regulation experiments, Western blots and antibodies

All siRNAs used to down-regulate expression of TAK1, TRAF6 or MEKK3 proteins were purchased from Origene, as well as siRNA No Target. The siRNAs were used at a final concentration of 10 nM. HEK293T-NF-κB cells were seeded in 24-well plate and were transfected the following day with siRNAs using the siTran 1.0 transfection reagent (Origene). The next day, culture medium was removed and replaced with complete medium. Eight hours after, cells were transfected with siRNAs alongside with plasmids encoding HIV-1 Env, HIV-2 Env or BST-2.

To perform knockdown of BST-2 in Jurkat-NF-κB cells, a TRC lentiviral human BST-2 shRNA vector (pLKO.1 vector) was obtained from Dharmacon (Clone ID: TRCN0000107015). We generated pseudotyped viruses to transduce Jurkat-NF-κB cells (a shRNA No Target was also used as a control for further experiments). Cellular clones of both Jurkat-NF-κB shRNA No Target and Jurkat-NF-κB shRNA BST-2 were selected by limiting dilutions in culture medium containing puromycin (2 μg/mL).

Expression levels of HIV-2 Env proteins or human TAK1, TRAF6, MEKK3 proteins were determined by Western blotting from HEK293T-NF-κB cell lysates used in the NF-κB activity experiments. The expression levels of human BST-2 were assessed in Jurkat-NF-κB cells. Samples were added to 2x Laemmli sample buffer, boiled for 5 minutes, separated by electrophoresis on 12% SDS-polyacrylamide gels, and transferred to nitrocellulose membranes. The membranes were blocked with 5% BSA in TBS containing 0.1% Tween-20 for 1 hour and probed overnight at 4°C with antiserum to HIV-2 ST gp120 (NIH AIDS Reagent Program, cat: 1410), or with mouse anti-TAK1, anti-TRAF6 or anti-MEKK3 antibodies (all from Origene). A rabbit monoclonal anti-BST-2 antibody (Abcam, cat: ab134061) was used to detect BST-2. Endogenous β-actin was detected with rabbit monoclonal anti-β-actin antibody (Cell Signaling, cat: 4970). Membranes were rinsed three times for 5 min in TBS 0.1% Tween-20 and blots were probed with either HRP-conjugated anti-rabbit or anti-mouse secondary antibody (Cell Signaling, cat: 7074 and 7076, respectively). Membranes were then rinsed three times and treated with the Amersham ECL Prime Western Blotting Detection Reagent (GE Healthcare Life Sciences), and blots were analyzed with the ChemiDoc XRS+ System (Bio-Rad).

### Quantitative RT-PCR

To determine mRNA levels of four selected genes targeted by the NF-κB transcription factor, Jurkat-NF-κB cells from remaining replicates of infection assays (24h and 48h post-infection) were harvested, washed twice in PBS and counted by Trypan blue exclusion method. Total RNAs were extracted using the RNeasy Plus Mini Kit (Qiagen) and were immediately subjected to reverse transcription with the Transcriptor First Strand cDNA Synthesis Kit (Roche). Each representative cDNAs of *il-6, il-21, ifn-β* and *bcl-2* genes were amplified with specific primers and quantified by qPCR using a LightCycler 96 system (Roche). The mRNA levels of *il-6, il-21, ifn-β* and *bcl-2* were compared to those of *gapdh* gene. The ratios were obtained using the 2^-ΔΔCT^ method.

Specific primers used in this study were as follows: IL-6: 5’-GAACTCCTTCTCCACAAGCGC-3’ and 5’-GTGGTTATTGCATCTAGATTCTTTGCC-3’; IL-21: 5’-GAGATCCAGTCCTGGCAACATG-3’ and 5’-CTTCACTTCCGTGTGTTCTAGAGG-3’; BCL-2: 5’-CACGCTGGGAGAACAGGGTAC-3’ and 5’-GTTGACGCTCTCCACACACATG-3’; IFN-β1: 5’-GACCAACAAGTGTCTCCTCC-3’ and 5’-GGCCTTCAGGTAATGCAG-3’; GAPDH: 5’-GAGTCAACGGATTTGGTCGTATTG-3’ and 5’-GCAGGAGGCATTGCTGAT GATC-3’ (all from Eurofins Genomics).

## AUTHOR CONTRIBUTIONS

Conceptualization, F.E.D. and B.K.M.; Formal analysis, F.E.D., G.D., M.L., J.R., L.T., M.S. and B.K.M.; Investigation and methodology, F.E.D., G.D., M.L.; Supervision, F.E.D.; Validation, F.E.D., G.D., M.L., K.S., E.N., J.R., L.T., M.S. and B.K.M.; Writing – original draft, F.E.D., G.D., M.L., K.S., E.N., J.R., L.T., M.S. and B.K.M. All authors have read and agreed to the published version of the manuscript.

## ACKNOWLEDGEMENTS

We thank Patrick Goubau (IREC Institute) and Thomas Michiels (de Duve Institute) for the fruitful discussions regarding this study and for their productive suggestions. The authors declare no conflict of interest regarding the publication of this manuscript.

## Notes

### Competing Interest Statement

The authors have declared no competing interest.

